# uvCLAP: a fast, non-radioactive method to identify *in vivo* targets of RNA-binding proteins

**DOI:** 10.1101/158410

**Authors:** Daniel Maticzka, Ibrahim Avsar Ilik, Tugce Aktas, Rolf Backofen, Asifa Akhtar

## Abstract

RNA-binding proteins (RBPs) play important and essential roles in eukaryotic gene expression regulating splicing, localization, translation and stability of mRNAs. Understanding the exact contribution of RBPs to gene regulation is crucial as many RBPs are frequently mis-regulated in several neurological diseases and certain cancers. While recently developed techniques provide binding sites of RBPs, they are labor-intensive and generally rely on radioactive labeling of RNA. With more than 1,000 RBPs in a human cell, it is imperative to develop easy, robust, reproducible and high-throughput methods to determine in vivo targets of RBPs. To address these issues we developed uvCLAP (UV crosslinking and affinity purification) as a robust, reproducible method to measure RNA-protein interactions *in vivo*. To test its performance and applicability we investigated binding of 15 RBPs from fly, mouse and human cells. We show that uvCLAP generates reliable and comparable data to other methods. Unexpectedly, our results show that despite their different subcellular localizations, STAR proteins (KHDRBS1-3, QKI) bind to a similar RNA motif *in vivo*. Consistently a point mutation (KHDRBS1^Y440F^) or a natural splice isoform (QKI-6) that changes the respective RBP subcellular localization, dramatically alters target selection without changing the targeted RNA motif. Combined with the knowledge that RBPs can compete and cooperate for binding sites, our data shows that compartmentalization of RBPs can be used as an elegant means to generate RNA target specificity.

## Introduction

Transcriptional control of gene expression is a highly regulated and intensely studied phenomenon that requires a plethora of DNA interacting proteins and other upstream factors that integrate intracellular and extracellular information to effect an appropriate transcriptional response (Kornberg, 1999; Vaquerizas et al., 2009). And yet, after a gene is fired, the message of the emerging RNA can be changed, muted, enhanced, localized or delayed through the local and global secondary structures the RNA forms and through the collective action of many RNA-binding (Gerstberger et al., 2014) and RNA-modifying proteins (Lewis et al., 2017) that act on these RNA transcripts via binding to specific RNA elements (Ray et al., 2013), or via changing the structure of the RNA (Kedde et al., 2010; Kenny and Ceman, 2016). Understanding the fate of an RNA not only requires a detailed understanding of its primary sequence and relative abundance but also what kind of RNP it is packaged into through the action of RNA helicases (Bourgeois et al., 2016) and other RNA-binding proteins (Gehring et al., 2017), both in the nucleus (Conrad et al., 2016) and in the cytoplasm (Oshiumi et al., 2016).

Several methods are available to identify binding sites of RNA-binding proteins on mRNAs and other RNAs. Low-resolution techniques such as RNA immunoprecipitation (RIP) (Niranjanakumari et al., 2002; Zhao et al., 2010) allow the identification of bound transcripts and are generally carried out after formaldehyde crosslinking to stabilize RNA-protein interactions and also to reduce potential in-solution re-binding artefacts (Mili and Steitz, 2004). Highresolution techniques for the identification of concise binding sites include HITS-CLIP (Licatalosi et al., 2008), iCLIP (König et al., 2010), PAR-CLIP (Hafner et al., 2010) and CRAC (Granneman et al., 2009) where protein-RNA interactions are permanently sealed by UV-crosslinking (Wagenmakers et al., 1980), which allows researchers to use RNases to trim the bound RNA, increasing resolution and specificity.

CLIP-seq approaches have become the principal experimental method for determining binding sites of RBPs. To date, CLIP-seq has been used to determine binding sites of hundreds of RBPs (Wang et al., 2015; Yang et al., 2015). These experiments are the basis of the recent advances in the elucidation of RBP-mediated regulation, offering unprecedented views of the complex nature of post-transcriptional regulation. For uncovering more complex relationships within the context of post-transcriptional regulation, however, a single CLIP-seq experiment will not be sufficient to determine the regulatory relationships in sufficient detail. In many cases this will mean increasing the experimental efforts. Researchers will need to probe multiple RBP isoforms (possibly differing in effect and localization), investigate RBPs with modified or disabled functionalities (e.g. variants lacking parts of RNA-binding motifs or helicase-dead mutants), examine multiple candidates suspected to be involved in a given regulatory process and also be able to compare results from multiple cell types and knockdown conditions. Widespread application of current CLIP-seq methods in this manner is thwarted by the high effort required --- the execution of a single experiment can take a week or more, the use of radioactivity makes the experimental handling cumbersome --- and the dependence on the availability of high-quality antibodies.

Because of the stringent washing procedure that can be employed due to crosslinking, CLIP-seq approaches have long been assumed to be mostly free of background. In consequence, experiments controlling for unspecific background have rarely been implemented. The first sequenced example of a CLIP-seq background control (a negative control immunoprecipitating a non-RNA-binding protein after crosslinking), however, found that nonspecific background was “common, reproducible and apparently universal among laboratories” (Friedersdorf and Keene, 2014). A comparison of 33 sets of PAR-CLIP sites with unspecific PAR-CLIP background showed an increase of background contamination for putative RBPs and experiments with small library sizes. Furthermore, sites supported by many reads were more frequently overlapped by background reads than sites supported by fewer reads. The mere occurrence of reads from a background control, however, is not sufficient to reject a site as nonspecific. Instead, the specificity of a site can only be reliably determined if the amount of background relative to the foreground signal is known. While similar amounts of foreground and background signal are indicative of a non-specific binding site that should be rejected, a low amount of background merely indicates high gene expression.

To address these challenges, we sought to develop a method that is fast, reliable, does not rely on radioactivity and that allows to quantify the amount of non-specific background to obtain transcriptome-wide high-resolution RNA-protein interaction maps with high specificity. To warrant the widespread utility of this method it should also be applicable to the three major model organisms Homo sapiens, Drosophila melanogaster and Mus musculus.

We performed uvCLAP experiments for a wide variety of RBPs, namely the KH domain-containing RBPs QKI-5, QKI-6, KHDRBS1-3 and hnRNPK, the DEAD-box helicases eIF4A1 and eIF4A3, the DExH-Box helicases MLE and DHX9, the member of the exon junction complex (EJC) MAGOH, and the mouse MSL complex members MSL1 and MSL2. In addition, we have probed mutant constructs of KHDRBS1, KHDRBS2 and MLE. In total, we have performed 23 uvCLAP experiments (investigating 15 RBPs) in human, mouse and fly.

## Results

The use of radioactive labeling of RNA that is covalently bound to a protein of interest has been used to verify the fact that a given protein interacts with RNA. Radioactive labeling has also been a hallmark of most CLIP-seq approaches. Since proteins tend to run at discrete positions on an SDS/LDS-PAGE setup, one can simply cut the labeled region and effectively remove contaminating RNA that migrates in the rest of the gel. In most cases, the protein-RNA complex is electrophoretically transferred to a nitrocellulose membrane, which typically (although not exclusively) binds to proteins (and consequently protein-RNA adducts) but not to free RNA, thus serving as another step for removing non-crosslinked RNA. We reasoned, that by employing a stringent tandem affinity purification protocol we could circumvent these time-consuming and critical steps and nonetheless remove contaminating free-RNA and proteins that might specifically or non-specifically co-purify with our protein of interest. To this end, we decided to use the HBH tag (Tagwerker et al., 2006) that allows rapid and ultra-clean purifications without the use of antibodies. In order to increase the versatility of this construct, we also added a 3xFLAG tag right before the HBH tag. We will refer this tag as the 3FHBH tag. An additional benefit of affinity purification using tagged proteins is the improved comparability across multiple conditions. Tagged constructs can be expected to have similar pulldown efficiency which would not be given when using antibodies.

We decided to include background controls employing non-specific pulldown of either tagged GFP or the expression vector carrying the 3FHBH tag without a gene insert. Furthermore, by preserving relative quantities even between different uvCLAP experiments, we can reliably determine the amount of background in our data.

After the cell lysates from RBP cell lines or control cell lines were homogenized, we made use of the tandem affinity tags with the order of polyhistidine pulldown followed by the streptavidin pulldown. The full protocol after the pulldowns lasts in total 4 days with the use of a single gel purification step (Figure 1A). We have validated the success of the specific protein pulldown by running the input (lane1 in Figure 1B), the imidazole eluate from the first polyhistidine purification (lane 2 in Figure 1B) and the final streptavidin eluate (lane 3 in Figure 1B) on a gel and by visualizing the bait with silver staining (Figure 1B). Various RBPs we have screened in human cells led to characteristic binding on their target RNA with a very weak and dispersed binding obtained from the background controls indicating the success of the tandem affinity pulldowns in removing the unspecific binding (Figure 1C).

**Figure 1.**
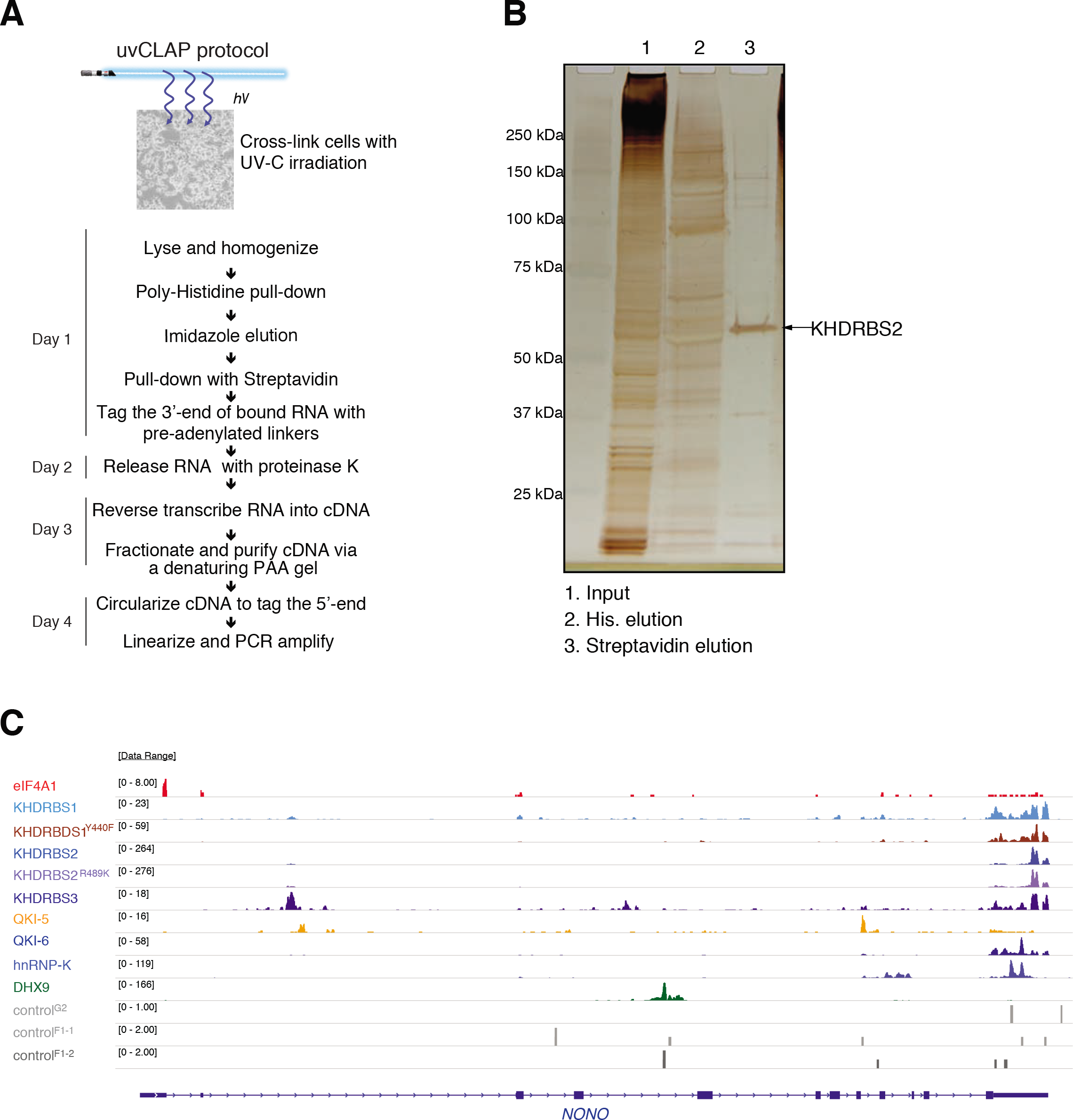
uvCLAP identifies in vivo targets of RBPs. **(A)** Experimental workflow of uvCLAP, starting from cells-on-plates to the generation of sequencing libraries. **(B)** Tandem affinity purification of 3xFLAG-HBH tagged KHDRBS2 under the highly-stringent uvCLAP conditions (see Methods). Lane 1: initial lysate, lane 2: eluate after the first step of purification, lane 3: final eluate from streptavidin beads. **(C)** IGV (Integrative Genomics Viewer) snapshot of uvCLAP profiles for eIF4A1, KHDRBS1, KHDRBS1^Y440F^, KHDRBS2, KHDRBS2^R489K^, KHDRBS3, QKI-5, QKI-6, hnRNPK and DHX9 on NONO gene. Biological replicates are merged for this representation, data range represents the coverage of uvCLAP reads. Only plus strand data is represented for clarity.

The controls allowed us to thoroughly evaluate the effects of the tandem purification. In more detail, we prepared multiplexed libraries containing both specific and unspecific pulldown eluates. Multiplexing was implemented using a triple-tag strategy that allowed us to distinguish pulldown conditions, biological replicates and size fractions. In addition, we used unique molecular identifiers (UMIs) for detecting individual crosslinked RNAs, i.e. crosslinking events, to improve the precision of our measurements. The multiplexed libraries were subjected to joint amplification and sequencing. In combination, this setup should allow to determine the relative quantities of crosslinked RNAs across multiplexed uvCLAP experiments (Figure 2A).

**Figure 2.**
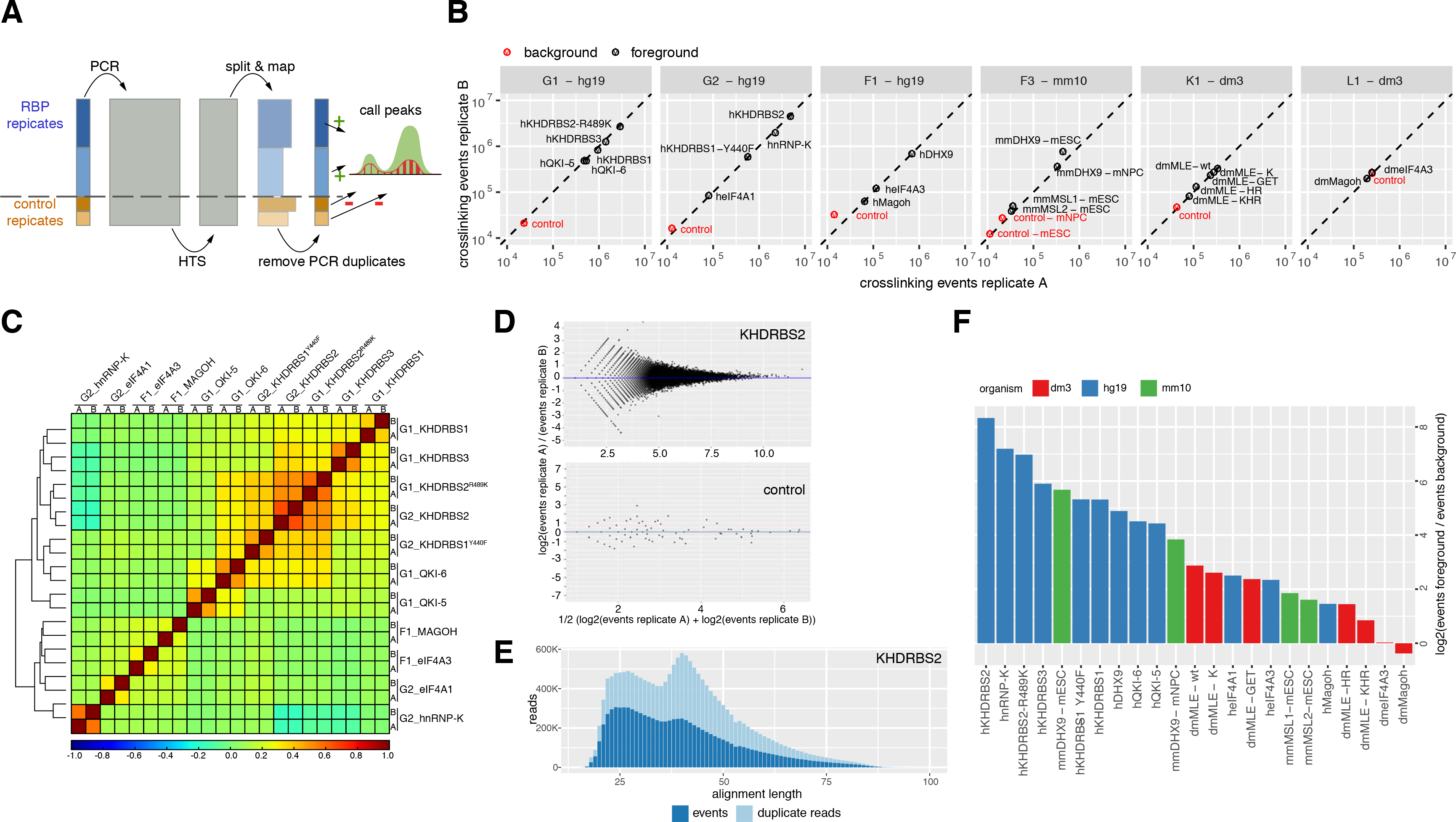
uvCLAP is a quantitative and reproducible assay. **(A)** Visualization of quantitative uvCLAP workflow. Replicates of pulldowns and controls are combined prior to PCR and jointly subjected to high-throughput sequencing. The resulting reads are split according to inline barcodes and barcodes are merged according to unique molecular identifiers (UMIs), recovering the relative RNA quantities present in the input libraries. **(B)** Comparison of the total number of crosslinking events identified for pairwise biological replicates of 23 pulldown conditions (black) and 7 unspecific controls (red). **(C)** Pairwise Spearman correlations of crosslinking events of uvCLAP replicates on merged JAMM peaks for human KHDRBS1-3, KHDRBS1^Y440F^, KHDRBS1^R489K^, QKI-5, QKI-6, MAGOH, eIF4A1, eIF4A3 and hnRNPK. **(D)** MA-plots comparing crosslinking event counts for pairwise biological replicates of KHDRBS2 and the corresponding background control for genomic 100 nucleotide bins covered by at least 2 crosslinking events in both replicates. The median log2 fold change is indicated in blue (see Supplementary Figure 2 for the full set of plots for all pulldown conditions). **(E)** Number of reads categorized as crosslinking events and PCR duplicates dependent on alignment length as proxy for cDNA insert size (see Supplementary Figure 3 for the full set of plots for all pulldown conditions). **(F)** Log2-ratios of crosslinking events to unspecific events from background controls for 23 pulldown conditions.

This setup allowed us to address the main challenges typically arising with the empirical measurement of CLIP-seq background. Stringent washing conditions typically result in small library sizes requiring high numbers of PCR cycles in order to be suitable for a dedicated sequencing run (Van Nostrand et al., 2016). In addition, when foreground and control are subjected to separate amplification and sequencing, the quantitative relationship between foreground and control remains unclear.

### uvCLAP determines the amount of crosslinked RNA with high precision

To determine to what extent the multiplexed treatment of specific and unspecific pulldown conditions would preserve the quantity of detected RNA fragments, we compared the total number of crosslinking events between pairwise biological replicates of 23 specific and 7 unspecific pulldown conditions in 6 multiplexed uvCLAP runs in Homo sapiens, Drosophila melanogaster and Mus musculus (Figure 2B, Supplementary Table 1), finding an excellent agreement of the absolute number of crosslinking events between paired replicates (Supplementary Figure 1B, Pearson correlation 0.997, n=30, p-value < 2.2e-16). This indicates that uvCLAP crosslinking events capture total library RNA counts exceptionally well.

We next asked, to what extent relative quantities were preserved at the binding site level. To this end, we compared biological replicates independently of the eventual peak calling on the level of genomic bins. For all 100 nucleotide bins covered by at least 2 crosslinking events in both replicates, we calculated the fold changes between the replicates (Figure 2D and Supplementary Figure 2). The resulting distributions of fold changes were consistently centered near 0, indicating a good between-replicate agreement. The amount of noise was low for bins with high crosslinking counts and increased steadily for bins with lower counts, a feature commonly observed with RNA-seq data (Anders and Huber, 2010).

We next asked, what library normalization, if any, would be necessary for uvCLAP. Library normalization is a method commonly used when comparing different samples, e.g. to determine differentially expressed genes or differential binding of RBPs, to compensate for differences in library preparation such as PCR amplification and sequencing depth. Normalisation methods employed for RNA-seq, e.g. the median ratios of observed counts (Anders and Huber, 2010), or CLIP-seq, e.g. the MA-plot normalization method used by dCLIP (Wang et al., 2014), however, are only feasible when a large number of bound regions --- ideally of similar binding strength --- are shared. For that reason, library normalisation for uvCLAP would only be feasible between biological replicates of the same protein, but not between uvCLAP foreground and background. Under certain conditions, however, the direct use of counts without further normalisation was shown to yield the best results for RNA-seq experiments (Li et al., 2015). We reasoned, that if relative quantities of detected RNAs were preserved by uvCLAP, we could forego library normalisation and directly use counts of crosslinked RNAs to determine enrichment over background of peaks. We determined library normalization factors for uvCLAP biological replicates using the median ratios of counts method (Anders and Huber, 2010; Robinson and Oshlack, 2010) (Supplementary Table 2). Normalization factors for 23 of the 30 pairwise replicates were 1, indicating that no library normalization would be required. The average normalization factor across all replicates was 0.96, indicating that uvCLAP counts accurately reflect crosslinked RNA quantities on the site level. In summary, we have shown that uvCLAP is suitable for both the quantification of total library RNA and local site counts within a multiplexed and jointly amplified uvCLAP library.

### Global enrichment over background is protein specific

Having established that uvCLAP crosslinking events accurately account for the amount of RNA in specific and unspecific libraries, we investigated whether we can see a clear difference between RBPs with large and small library sizes. For that purpose, we calculated the total enrichment of specific versus unspecific conditions based on the total number of crosslinking events per library (replicates combined) (Figure 2F, Supplementary Table 3) for different RBPs. This revealed that the library-wide enrichment of molecules of specific versus unspecific pulldown conditions can vary over three orders of magnitude. At the high end, we observed enrichment over background of more than 300-fold enrichment for human KHDRBS2, a large library with 9.3 million crosslinking events. At the low end, we found only a very slight enrichment of Drosophila eIF4A3 over background (foreground/background ratio 1.02) and no enrichment for Drosophila MAGOH (foreground/background ratio 0.77), two small libraries with 517,000 and 392,000 crosslinking events. This data confirms the observation of Friedersdorf and Keene, that background is especially pronounced for RBPs with small library sizes (Friedersdorf and Keene, 2014). These results also show that the assumption of the high specificity of CLIP-seq only holds under ideal conditions, i.e. for proteins that crosslink well and bind to a large number of sites. We conclude, that the quantification of background signal of a given CLIP-seq library is a requirement to reliably determine binding of the large number of putative RBPs not yet probed using CLIP-seq. Additionally, global enrichment over background can serve as a straightforward means to identify problematic experiments.

### cDNA-length specific amplification bias is mitigated by UMIs

Having separately tagged size fractions available, we observed that events from high size (H, cut around 75 nt) and mid size (M, cut around 50 nt) size fractions were represented by more duplicate reads than the short size (L, cut around 25 nt) fractions which may indicate a length bias introduced by PCR amplification (Dabney and Meyer, 2012). This effect was very pronounced for the H libraries, to the extent that despite large numbers of reads only few crosslinking events could be identified. In consequence, we decided not to use the H size fractions and omitted their sequencing for Drosophila eIF4A3 and MAGOH uvCLAP.

To further investigate the influence of cDNA length on the amount of duplicate sequences, we evaluated the number of uniquely aligned reads and resulting events of M- and L-fraction in relation to the lengths of the corresponding alignments (Figure 2E and Supplementary Figure 3). The resulting distributions frequently revealed an excess of PCR duplicates for cDNAs in the range of 40-60 nts. These seem to have been amplified more efficiently than shorter or longer cDNAs, leading to an excess of these lengths. This bias largely disappeared at the level of crosslinking events, leading to mostly unbiased distributions over the full range of cDNA sizes. In consequence uvCLAP maintains a wide range of cDNA lengths, suggested by (Haberman et al., 2017) to avoid cDNA-end constraints that may hinder accurate identification of bound sites.

### Peak calling analysis for uvCLAP data

Peak calling refers to the identification of discrete binding sites from reads or binding events. Usually peak calling also includes filtering of identified peaks in order to reduce the number of false positive binding sites. However, existing strategies such as the presence of specific mutations caused by crosslinked nucleotides (Corcoran et al., 2011; Zhang and Darnell, 2011), enrichment over shuffled input signal (Althammer et al., 2011; Xue et al., 2009; Yeo et al., 2009), and modelling of read-count distributions (Chen et al., 2014; Uren et al., 2012) could not be used due to the specifics of our data.

The basic idea of the most widely used peak calling methods is to compare the number of crosslinking events of a region to a background distribution in order to determine regions with significantly enriched number of events. There are in general two principal approaches for determining a background distribution. The first approach, namely shuffling of the gene-wise input signal, assumes a uniform distribution for the background. However, this uniformity is neither observed with uvCLAP nor PAR-CLIP background (Friedersdorf and Keene, 2014). The second approach uses global read-count distributions and assumes that most reads constitute background. This assumption is again not indicated by our data. A third possibility, assuming that our libraries are mostly free of background, would be to omit any filtering after initial peak identification (König et al., 2010; Licatalosi et al., 2008). This was also rejected as there is varying amount of unspecific binding seen in our data (König et al., 2010; Licatalosi et al., 2008).

Thus, we looked for a peak calling method that is able to incorporate quantitative background samples into the peak calling procedure. These requirements are satisfied by JAMM (Ibrahim et al., 2015), a universal peakfinder, which is able to integrate information from uvCLAP biological replicates and background controls. JAMM first determines genomic bins likely to contain binding sites based on enrichment of foreground over background signal and the signal-to-noise ratio of a bin compared to the signal-to-noise ratio over the whole chromosome. Peak boundaries are then determined for all bins that satisfy these constraints for all replicates. JAMM typically reports a large number of peaks, allowing for flexible further filtering. We reasoned that this approach would give us the most flexibility and help to give an accurate view of the uvCLAP data. The lowest number of JAMM peaks was obtained for mmMSL1 (479 peaks), the highest number of peaks resulted for KHDRBS2 (298,135 peaks) (see Supplementary Table 4).

We used the JAMM peaks to calculate pairwise correlations of the number of crosslinking events located on merged peaks for the biological replicates of human proteins KHDRBS1-3, KHDRBS1^Y440F^, KHDRBS2^R489K^, QKI-5, QKI-6, eIF4A1, eIF4A3, MAGOH and hnRNPK (Figure 2C). Correlations between biological replicates were in all cases higher than correlations to replicates of different pulldown conditions. The average correlation between biological replicates of 0.46 was much increased over the average correlation of 0.08 between unrelated replicates (excluding comparisons between KHDRBS1-3 and between QKI-5 and QKI-6). We also observed high correlations between replicates of KHDRBS2 and its pseudo-mutant construct KHDRBS2^R489K^ but not between replicates of KHDRBS1 and its mutant KHDRBS1^Y440F^ that differ in cellular localization (see below for a detailed discussion of the mutant constructs).

### uvCLAP background is distinct from foreground

To obtain a high-level view on uvCLAP background, we determined the distributions of control crosslinking events on different classes of genes (Supplementary Figure 1C-E). Here, the two fly control libraries exhibited similar distributions over the targeted classes of genes, the most prominent class of targets being ribosomal RNAs. The two mouse libraries also showed similar distributions, many events were located on intronic and intergenic regions. The enrichment of ribosomal RNAs found for fly background was not observed for human and mouse controls. The dominant location of background was introns in human datasets. The background reads from the three human libraries (G1, G2 and F1) had comparable occurrences of events on ribosomal, noncoding RNAs and pseudogenes. We observed a larger variability of the distributions on regions of coding genes, intergenic and antisense regions than observed for fly and mouse backgrounds (Supplementary Figure 1C compared to 1D and 1E). The control sample of library F1 was enriched for intronic, antisense and intergenic locations, while the control for library G1 showed increased 3’-UTR targeting compared to F1 and G2 (Supplementary Figure 1C). This variability may be indicative of the stochastic nature of background events, however, we cannot exclude the possibility of a weak shadowing of one of the foreground samples in the background library.

We next evaluated the connection between uvCLAP foreground to background signal by determining the number of JAMM peaks more than 50 nucleotides apart from any read of the corresponding control (Supplementary Table 4). 94.15% of peaks derived for the 23 pulldown conditions were not located in the vicinity of control reads; 16 of 23 pulldown conditions had more than 80% of peaks without evidence of surrounding control reads. Proteins with higher correspondence to control reads are known to inefficiently crosslink (hsMAGOH), had low numbers of binding sites (mmMSL1 and mmMSL2) or were mutant constructs with one or two disabled double-stranded RNA-binding domains (dmMLE-HR and dmMLE-KHR). The largest coincidence with the controls was observed for dmeIF4A3 and dmMAGOH with 26.21% and 16.39% of peaks in the vicinity of control reads.

The overwhelming majority of sites detected by uvCLAP were independent of background. The seven proteins that most strongly coincided with of overlap with background signal also had the lowest global foreground to background ratios (below 5). Thus, we conclude, that quantitative uvCLAP background can serve as a straightforward means to estimate the quality of a uvCLAP experiment.

### uvCLAP recovers known binding preferences of DEAD-box helicases eIF4A1 and eIF4A3

We decided to benchmark our protocol with two well-known and highly related DEAD-box helicases eIF4A1 and eIF4A3. We chose these proteins for two reasons: first, both proteins have well-defined RNA-binding behaviour. eIF4A1 is part of the cytoplasmic eIF4 complex that scans the 5’-UTRs of mRNA in search of a start codon, thus we anticipated a strong 5’UTR enrichment as a proof that uvCLAP works. eIF4A3, on the other hand, is a part of the predominantly nuclear exon junction complex (EJC) that binds to 20-30nt upstream of exon junctions, thus we expected to detect an exonic enrichment and a positional bias for eIF4A3. Second, especially for eIF4A3, it is clear that this protein interacts mainly with the sugar-phosphate backbone of the target RNA, which makes it a poor UV-crosslinking protein, and hence a challenging protein to use as a benchmark (Singh et al., 2014).

As expected, 5’-UTRs were the most abundant type of eIF4A1 targets (Figure 3H), in total 63% of the 2,653 eIF4A1 peaks located on protein-coding gene regions were annotated as 5’-UTRs. For human eIF4A3 and MAGOH, coding exons were the most abundant type of targets (Figure 4A), with 78% of hseIF4A3 and 81% of hsMAGOH peaks located on proteincoding genes targeting regions annotated as coding exons.

**Figure 3.**
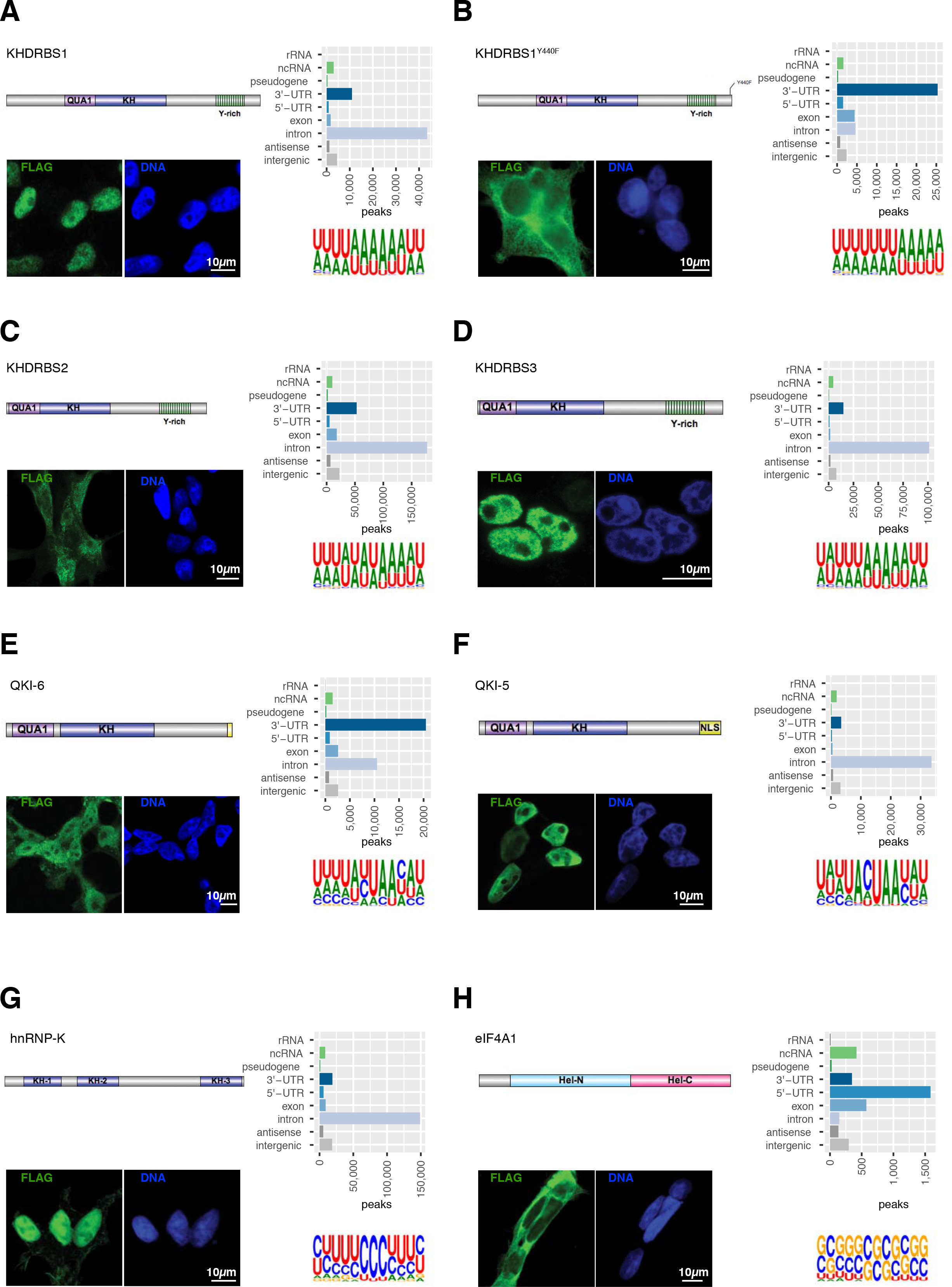
Cellular localization of RBPs do not change their preferred RNA motif. Cellular localization of FLAG-tagged RBPs; number of JAMM peaks located on genomic target classes with each peak assigned once to the category with highest priority corresponding to the order rRNA, ncRNA, pseudogene, 3’-UTR, 5’-UTR, exon, intron, antisense, intergenic (see Supplementary Table 5 for the counts of peaks on genomic targets); and Sequence motifs determined by GraphProt (Maticzka et al., 2014) for human proteins **(A)** KHDRBS1, **(B)** KHDRBS1^Y440F^, **(C)** KHDRBS2, **(D)** KHDRBS3, **(E)** QKI-6, **(F)** QKI-5, **(G)** hnRNPK, and **(H)** eIF4A3.

**Figure 4.**
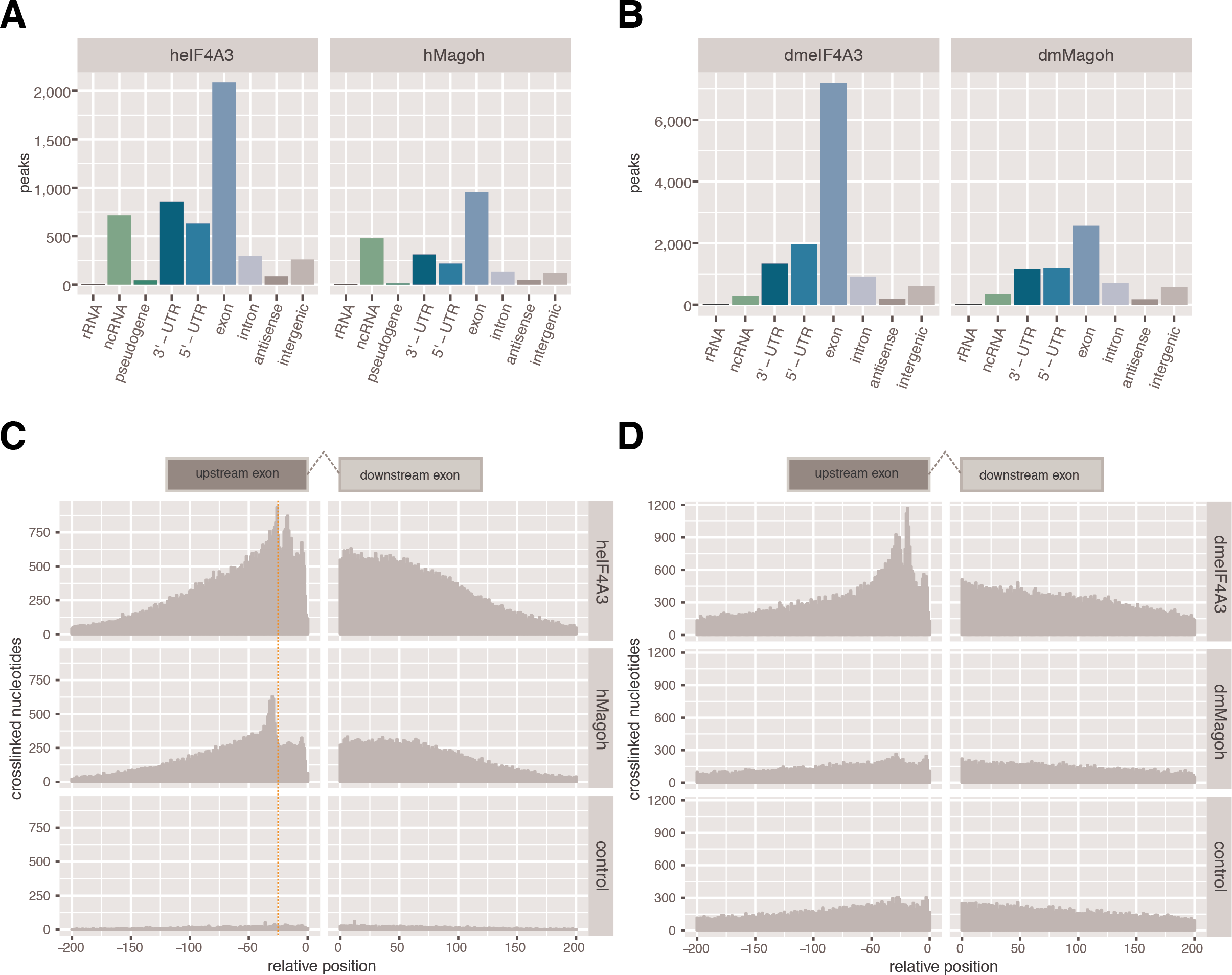
eIF4A3 binds upstream of exon-exon junctions. **(A)** Number of JAMM peaks located on genomic target classes for human eIF4A3 and MAGOH. Priority of target classes as in Figure 3. **(B)** Number of JAMM peaks located on genomic target classes for Drosophila eIF4A3 and MAGOH. Peaks were assigned according to the order rRNA, ncRNA, 3’-UTR, 5’-UTR, exon, intron, antisense, intergenic. **(C)** Histogram of crosslinked nucleotide positions for human eIF4A3, MAGOH and the corresponding control. eIF4A3 and MAGOH show increased binding 20-30 nucleotides upstream of exon-exon junctions, binding decreased with increasing distance to exon-exon junctions whereas the control is uniformly distributed. **(D)** Histogram of crosslinked nucleotide positions for Drosophila eIF4A3, MAGOH and the corresponding control. eIF4A3 shows increased binding 20-30 nucleotides upstream of exon-exon junctions whereas crosslinked nucleotides of MAGOH and the control are not specifically enriched.

To accurately determine the binding of human and fly EJC (eIF4A3 and MAGOH) near exon-exon junctions, we realigned the libraries using the splice-aware mapper hisat2 (Kim et al., 2015; Pertea et al., 2016) (version 2.0.5, parameters --fr --no-mixed --no-discordant) and otherwise performed the same processing as done for the other uvCLAP sets and determined the locations of crosslinked nucleotides relative to 5’- and 3’-exon ends. We observed an enrichment of crosslinked nucleotides for human eIF4A3 near the expected positions upstream of exon-exon junctions (Figure 4C). Differently to the findings reported by Sauliere and colleagues (Sauliere et al., 2012), binding detected by uvCLAP outside of canonical positions was not uniform in the region 200nt upstream of exon 3’-ends, but diminished with increasing distance to exon-exon junctions. In contrast, crosslinked nucleotide positions of the background control were uniformly distributed in relation to exon-exon junctions. For MAGOH we observed a similar positional pattern that was shifted slightly upstream to that of eIF4A3.

We obtained very similar results for Drosophila eIF4A3 (dmeIF4A3). 68% of the peaks located on protein coding genes were annotated as coding exons (Figure 4B). We also observed a positional pattern similar to hseIF4A3 (Figure 4D). In contrast to dmeIF4A3, Drosophila MAGOH peaks were only slightly enriched for coding exons. The corresponding profile relative to exon-exon junctions was quite uniform in the regions 200 nucleotides up- and down-stream of exon-exon junctions and very similar to that of the background control, confirming our initial expectation based on the foreground-to-background ratio and high overlap with background that we could not detect binding of this protein.

In summary, uvCLAP recovered the known position-specific binding of DEAD-box helicase eIF4A1 on 5’-UTRs and of the EJC members eIF4A3 and MAGOH near exon junctions. The results for Drosophila MAGOH highlight the usefulness of quantitative background controls for assessing the quality of experimental data at an early stage of the analysis, which should be especially helpful when evaluating uvCLAP experiments for RBPs with unknown binding preferences.

### uvCLAP recovers binding preferences of KH-domain containing proteins QKI, KHDRBS1, KHDRBS2, KHDRBS3 and hnRNPK

Next, we decided to evaluate QKI, a protein with a previously identified sequence-specificity and is known to regulate target-mRNA stability (Teplova et al., 2013) and circRNA formation (Conn et al., 2015). QKI has multiple isoforms that are slightly different at the C-termini. Despite being small, these differences have a profound effect on the subcellular localization of QKI: the shorter isoforms lack a nuclear localization signal that makes them predominantly cytoplasmic (Pilotte et al., 2001). This behaviour is at odds with QKI biology as it is highly related to the primary branch point recognizing protein SF1. Indeed, the QKI consensus motifs derived from SELEX (Galarneau and Richard, 2005) and PAR-CLIP (Hafner et al., 2010) are very reminiscent of the branch point sequence.

We chose to evaluate the predominantly nuclear isoform QKI-5 as well as the shorter isoform QKI-6 that is reported to be both nuclear and cytoplasmic (Conn et al., 2015). We could confirm this behaviour for our triple-tagged constructs via anti-FLAG staining (Figure 3E,F). In accord with this observation, peaks of the nuclear isoform QKI-5 were located predominantly on intronic regions whereas peaks of QKI-6 were located to a large extent on 3’-UTRs as well as intronic regions (Figure 3E,F). The GraphProt (Maticzka et al., 2014) motif for QKI-5 closely resembled the known QKI core motif ACUAAY (Galarneau and Richard, 2005; Ryder and Williamson, 2004), the motif for QKI-6 better resembled the consensus motif AYUAAY identified from highly expressed PAR-CLIP read clusters (Hafner et al., 2010) (Figure 3E,F).

We next compared uvCLAP QKI binding sites with sites derived from QKI PAR-CLIP (Hafner et al., 2010) (retrieved from CLIPdb (Yang et al., 2015), peaks called by PARalyzer (Corcoran et al., 2011)). JAMM produces two sets of peaks, a full set of peaks containing also small peaks and peaks with few reads, and a smaller, filtered set where these peaks are removed (in the following named JAMM full and filtered, respectively). To facilitate the comparison of the peaks from the two methods and also integrate small peaks, all peaks were extended to a minimal length of 41 nucleotides and overlapping or abutting peaks were merged.

A comparison of the overlaps between the full uvCLAP peak sets identified 7,761 sites shared between QKI-5 and QKI-6 and 2,685 sites shared between uvCLAP and PAR-CLIP, leaving several thousand sites exclusive to each of the three experiments (Figure 5A, Supplementary Figure 4A). Using the filtered uvCLAP peaks, the number of sites shared between QKI-5 and QKI-6 uvCLAP was reduced to 30.49% (2.336 sites) compared to the full set, but also reduced the total number of uvCLAP sites to 32.37% for QKI-5 (13,530 sites) and to 34.23% for QKI-6 (12,832 sites) while the number of sites shared between uvCLAP and PAR-CLIP was reduced to 47.78% (1,283 sites) compared to the full list of peaks. Possible reasons for the low overlap between uvCLAP and PAR-CLIP sites are mainly technical differences between the two methods. Those include differences in RNase digestion, the wavelength used for crosslinking, preferential crosslinking to 4-thiouridine (Kishore et al., 2011) as well as the use of different isoforms and differently tagged expression vectors.

**Figure 5.**
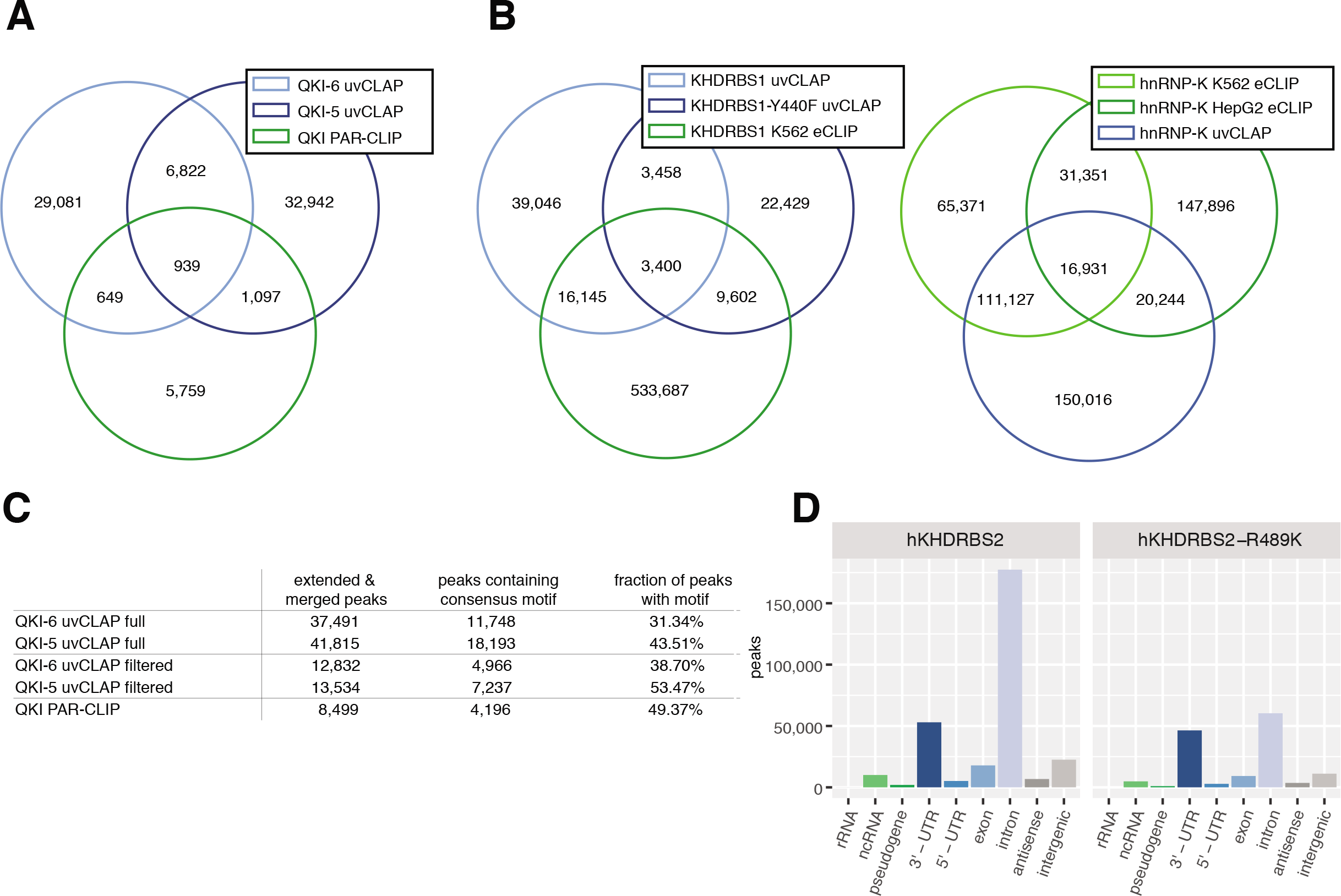
Comparison of uvCLAP to other methods. **(A)** Overlap between the full sets of QKI-5 and QKI-6 uvCLAP and QKI PAR-CLIP peaks. Peaks were extended to a minimum length of 41 nt, overlapping or abutting sites were merged. **(B)** (left) Overlap between the full sets of KHDRBS1, KHDRBS1-Y440F uvCLAP and KHDRBS1 K563 eCLIP peaks. (right) Overlap between the full sets of hnRNPK uvCLAP, hnRNPK K562 eCLIP and hnRNPK HepG2 eCLIP peaks. Peaks were extended to a minimum length of 41 nt, overlapping or abutting sites were merged. **(C)** QKI-5, QKI-6 and QKI PAR-CLIP sites containing the consensus motif AYUAAY. **(D)** Number of JAMM peaks located on genomic target classes. Each peak was assigned once to the category with highest priority corresponding to the order rRNA, ncRNA, pseudogene, 3’-UTR, 5’-UTR, exon, intron, antisense, intergenic.

To further investigate the quality of uvCLAP peaks, we determined the occurrence of the consensus motif AYUAAY (Figure 5C). For the full sets of uvCLAP peaks, the fraction of peaks harboring the consensus motif was smaller for uvCLAP (QKI-6: 31.34%, QKI-5: 43.51%, QKI PAR-CLIP 49.37%), however, uvCLAP identified a larger number of sites harboring the consensus motif (QKI-6: 12,212, QKI-5: 19,021) compared to 4,209 sites identified by PAR-CLIP. In total, uvCLAP identified 26,503 sites containing the consensus motif, of which 24,939 were exclusive to uvCLAP. Use of the filtered uvCLAP sets increased the fraction of peaks containing the consensus motif (QKI-6: +7.36%, QKI-5: +9.96%), lead to a reduction of the number of peaks containing the motif by 57.73% (QKI-6) and 60,22% (QKI-5), but still identified 9,992 peaks containing the consensus motif that were not found by PAR-CLIP (Figure 5C). In summary, we were able to recover known QKI sequence binding preference in uvCLAP data for two different QKI isoforms. Thus, uvCLAP identified several thousand novel QKI binding sites harboring the known consensus motif, which were not detected by PAR-CLIP.

Next, we evaluated three KH-domain containing RBPs KHDRBS1 (Matter et al., 2002), KHDRBS2 (Haegebarth et al., 2004; Wang et al., 2002) and KHDRBS3 (Haegebarth et al., 2004). We found that KHDRBS1, also known as Sam68, is nuclear as reported earlier (Matter et al., 2002) and to interact primarily with introns (Figure 3A). Similar to KHDRBS1, KHDRBS3 is also nuclear and it interacts mainly with introns (Figure 3D). On the other hand, we found that KHDRBS2 in our cells is mainly cytoplasmic and interacts with introns and 3’-UTRs (Figure 3C). Interestingly, all three proteins recognize a similar AU-rich RNA sequence (see below), irrespective of their subcellular localization (Figure 3A,C,D). We then cloned a point mutant of KHDRBS1, KHDRBS1^Y440F^ that was reported to disrupt a functional nuclear localization signal (NLS) (Lukong et al., 2005). KHDRBS1^Y440F^, as reported before, is indeed cytoplasmic (Figure 3B), and recognizes an AU-rich RNA sequence very similar to the wild-type KHDRBS1 as well as KHDRBS2 and KHDRBS3 but mostly at 3’-UTRs of mRNAs rather than introns (Figure 3B). Finally, we re-cloned KHDRBS2 with a serendipitous mutation R489K and re-created a cell line that expresses this variant. We found that KHDRBS2 and KHDRBS2^R489K^ have virtually identical RNA-binding profiles (Figure 1C), and correlate with each other extremely well (Figure 2C) with KHDRBS2 recognizing more intronic peaks compared to KHDRBS2^R489K^ (Figure 5D). Motif analysis of KHDRBS1-3 and the two point mutants revealed similar AU-rich motifs for all proteins (Figure 3A-D), matching the known affinity of KHDRBS1 to UAAA (Lin et al., 1997) and U(U/A)AA repeats (Galarneau and Richard, 2005). A preference for UAA was also shown for all three KHDRBS proteins by RNAcompete (Ray et al., 2013). For KHDRBS1, 31.5% (19,545 sites) from the full set of peaks and 40.75% (7,902 sites) from the filtered set of peaks were also found in KHDRBS1 eCLIP for K563 cells (Van Nostrand et al., 2016) (Figure 5B, Supplementary Figure 4B). These results show that uvCLAP protocol, and the analysis pipeline described here, yields robust results even when independently generated cell-lines are used to probe the same or very similar proteins.

Since STAR proteins contain a KH-domain that acts as the RNA-binding module, we decided to clone and determine the binding sites and preferences of the prototypical KH-domain-containing protein hnRNPK. In our system hnRNPK appears to be nuclear (Figure 3G). Our uvCLAP data show that hnRNPK interacts mainly with cytosine-rich RNA, specifically when a RNA contains three or more consecutive cytosines (Figure 3G). Comparison with two hnRNPK eCLIP experiments in K562 and HepG2 cells revealed that 49.74% (148,480) of uvCLAP sites were matched by sites from one of the two eCLIP experiments. For the filtered list of uvCLAP peaks, the overlap dropped to 33.23% (21,281) of uvCLAP peaks (Figure 3G, Supplementary Figure 4C). The GraphProt motif determined from hnRNPK uvCLAP data (Figure 3G) closely resembled the consensus motif determined by SELEX (Thisted et al., 2001). Taken together our results show that KH-domains are not restricted to AU-rich motifs (KHDRBS proteins) but can also recognize C-rich sequences (hnRNPK).

## Discussion

uvCLAP shows great promise in understanding, identifying and characterizing targets of RNA-binding proteins *in vivo* without having to resort to labor-intensive techniques that use radioactive substances. Using highly-stringent buffers, we are able to wash-away nonspecifically interacting RNA and other non-covalent interaction partners. Straightforward multiplexing of experiments leads to further time savings by allowing to process multiple samples in parallel. We show that uvCLAP works well for human, mouse and fly cell lines.

A major benefit of joint amplification and sequencing of multiplexed uvCLAP experiments is the preservation of relative RNA quantities within the multiplexed library both at the global and at the binding site level. This allows the direct comparison of different experimental conditions without additional library normalization. While methods that necessitate library normalization only allow the comparison of conditions that share a large number of binding sites of similar strengths (Wang et al., 2014), uvCLAP allows the comparison of arbitrary conditions.

This unique feature allowed us to determine the actual extent of background contamination of uvCLAP libraries via comparison to controls using unspecific pulldown conditions. For most experiments, we observed large global enrichments of foreground over background events. In addition, the vast majority of called peaks were not located in the vicinity of background events, emphasizing the capability of uvCLAP tandem purification to effectively remove unspecific background. Experiments with lower global enrichment over the background controls and increased occurrence of background events near binding sites were either known to inefficiently crosslink, bind only to few sites, or were deliberately modified for impaired binding. This indicates that quantitative uvCLAP background controls are a good means to determine the quality of a given experiment. Moreover, preservation of relative RNA quantities makes uvCLAP ideally suited for the investigation of differential binding for investigating multiple protein isoforms or knockdown conditions. The straightforward comparability of arbitrary pulldown conditions also allows the detailed investigation of altered binding exhibited by RBP deficiency mutants.

Our results show that uvCLAP accurately identifies expected targets when used with proteins with well-defined functions such as eIF4A1 and eIF4A3 (this study) as well as MLE (accompanying manuscript) and DHX9 (Aktaş et al., 2017). In addition, we used uvCLAP to systematically identify the *in vivo* targets of several STAR proteins including a point mutant and a splice-variant that change the subcellular localization of the target protein. To our surprise, irrespective of their subcellular localization, KHDRBS1, KHDRBS1^Y440F^, KHDRBS2, KHDRBS2^R489K^ and KHDRBS3, all recognize a similar AU-rich motif. Furthermore, a highly related protein, QKI, both the long and short isoforms, recognize a branch-point-like sequence, ACUAA that is only different from the KHDRBS-motif by the frequent presence of a cytosine, showing the sensitivity of the uvCLAP analysis. Interestingly, hnRNPK, another KH-domain containing RBP, recognizes a CU-rich motif with central, consecutive cytosines, very similar to the *in vitro* determined binding motif (Thisted et al., 2001). These results collectively show that the KH domain can recognize an array of related RNA motifs, and KHDRBS proteins likely regulate a common set of mRNAs through similar binding sites that can be present in introns and/or 3’-UTRs of targets (Traunmüller et al., 2014).

The fact that, despite their different subcellular localizations, STAR proteins bind to similar RNA motifs and that we can identify distinct RNA motifs for highly related proteins, shows the versatility and robustness of uvCLAP. Thus, mapping RNA-protein interactions *in vivo* reveals the multi-faceted nature of RNA-binding proteins and shows that compartmentalization of RBPs can be used as a mechanism to generate RNA target specificity.

## Limitations

In order to minimize non-specific contaminants that arise from abundant RNA-species such as ribosomal RNAs, snRNA and tRNAs, uvCLAP relies on *in vivo* target protein biotinylation and a quick tandem-affinity purification under very stringent conditions. We achieved biotinylation either by exogenously expressing proteins of interest with a biotinylatable peptide or via introduction of the tag into the endogenous locus by CRISRP/Cas9. Due to this dependence on biotinylation, uvCLAP cannot be used when these methods are not feasible (e.g. primary cells) or when affinity-tagging changes target protein localization and/or activity.

## Acknowledgements

We thank U. Bönisch and the members of the sequencing facility of the Max Planck Institute of Immunobiology and Epigenetics. This work was supported by CRC992 (AA, RB), CRC 746 (AA) and CRC1140 (AA).

## Data Accession

The uvCLAP data in this study has been deposited to the Gene Expression Omnibus database under the accession numbers GSE87792 (MLE) and GSE85155 (all other proteins).

The reviewer links are as follows:

https://www.ncbi.nlm.nih.gov/geo/query/acc.cgi?token=yiuxysswvhsrlyf&acc=GSE87792

https://www.ncbi.nlm.nih.gov/geo/query/acc.cgi?acc=GSE85155

## Methods

### Cell culture and generation of stable cell lines

FLPin Trex HEK293 cells are maintained with DMEM-Glutamax (Gibco 31966) and 10% FBS. The original cell line was maintained in zeocin-and blasticidin-containing medium according to manufacturer’s protocol (Thermo Fisher Scientific Catalog no. R780-07) and the zeocin selection is exchanged with hygromycin upon transgene transfection for stable cell line generation. All the transgenes were cloned into pCDNA5-FRT/To (Thermo Fisher Scientific Catalog no. V6520-20) with a C-term 3xFLAG-HBH tag and were co-transfected with pOG44 plasmid with a 1:9 DNA concentration ratio as suggested by the manufacturer’s protocol. Cells were re-plated in different dilutions (1:2, 1:3 and 1:6) 24 hours after the transfection and selection with 150μg/ml hygromycin was initiated 48 hours after transfection. Cell lines were maintained with blasticidin and hygromycin at all times and the transgenes were induced with with 0.1 μg/ml doxycycline for 16 hours both for the uvCLAP and Immunofluorescence experiments.

Mouse embryonic stem cells (ESCs) were maintained with 15% FBS, 2000U/ml LIF, sodium pyruvate, NEAA, and 0.1nM beta mercaptoethanol supplemented into DMEM-Glutamax.

CRISPR/Cas9 facilitated endogenous tagging of the mouse Msl1, Msl2 and Dhx9 was performed in a mouse ES cell line (WT26 male ES cell line was a kind gift of Jenuwein Lab) as described earlier in (Aktaş et al., 2017).

### Immunofluorescence

Doxycycline induced cells were crosslinked with 4% methanol-free formaldehyde in PBS at room temperature for 10 minutes and permeabilized with 0.1% Triton-X and 1% BSA in PBS for 30 minutes at room temperature. Primary FLAG-M2 antibody was diluted (1:500) in PBS with 0.1% Triton-X and 1% BSA and incubated with fixed cells at 4°C for ∼16hrs. Fluorescently labeled secondary antibodies with the appropriate serotype were used reveal target proteins. Hoechst 33342 to stain DNA. Imaging was performed with a Leica SP5 confocal microscope.

### uvCLAP procedure

Doxycycline induced FLPin Trex HEK293 cells are rinsed with PBS and crosslinked with 0.15mJ/cm^2^ UV-C light. Following crosslinking, cells are pelleted by centrifugation, snap-frozen in liquid nitrogen and kept at −80°C until use. Cells are then defrozen on ice and lysed with 0.5mL of lysis buffer (FLAG immunoprecipitations: 50mM Tris.Cl, pH 7.4; 140mM NaCl, 1mM EDTA, 1% Igepal CA-630, 0.1% SDS, 0.1% DOC; His-pulldowns: 1X PBS, 0.3M NaCl, 1% Triton-X, 0.1% Tween-20), mildly sonicated and immunoprecipitated with anti-FLAG beads for 1hr at °C or incubated with His-Tag Pulldown Dynabeads (10103D, Thermo Fisher Scientific) for 10 minutes. The beads are washed with lysis buffer and bound material is eluted with 3xFLAG peptide (250μg/mL) or 250mM imidazole in respective lysis buffer. The eluate is then incubated with MyONEC1 beads to collect biotinylated target protein, after which the beads are washed with high-stringency buffers (0.1%SDS, 1M NaCl, 0.5% LiDS, 0.5M LiCl and 1% SDS, 0.5M LiCl) to aggressively remove non-specific interactors. 3’-linkers are then ligated with T4 RNA Ligase 1, excess adapters are washed away and 3’-tagged, cross-linked RNA is released with proteinase K digestion and column purification (Zymo DNA Clean & Concentrator). Reverse transcription is carried out with SuperScript III and barcoded reverse-transcription primers. After reverse transcription, relevant samples are mixed and the cDNA is separated on a 6% 6M Urea PAA gel. Size fractionated cDNA is then processed essentially as described in the iCLIP protocol to generate sequencing libraries.

### Silver gel staining

Silver Gel staining was performed using the Silver Quest Silver Staining Kit (Thermo Fisher LC6070).

### uvCLAP tri-barcode approach

The uvCLAP tri-barcode approach --- based on ScriptSeq PCR primers and two custom 5 nucleotide barcodes adjoining the 5’- and 3’-adapters --- allows to flexibly tag source libraries according to multiple experimental conditions. This tagging strategy enabled us to tag pulldown conditions, size fractions and biological replicates prior to PCR amplification and sequencing. Custom 5’-tags and 3’-tags were used to distinguish pulldown condition and biological replicates. Size fractions were differentiated using commercially available ScriptSeq PCR primers.

To ensure optimal in-silico separability of experimental conditions, we used edittag (Faircloth and Glenn, 2012) to design tags robust to indel (insertion/deletion) and substitution errors. For this purpose, we created a set of 5 nucleotide tags with minimal pairwise edit-distance of 3, ensuring that 1 indel or substitution error can be corrected. The initial set of 31 candidate tags, created so as not to contain polybases or self-complements, was further filtered for tags containing nucleotide repetitions at either end (11 tags) and tags reverse complementary to the adapter (5 tags), leaving 15 tags for use by uvCLAP (Supplementary Table 6). The combination of tags used for each multiplexed library was chosen to provide at least one nucleotide detected by the red and green color channels used by Illumina sequencers at each position.

To distinguish pairwise biological replicates, we created semi-random tags according to patterns DRYYR and DYRRY (IUPAC ambiguity code, D: not C, R: purine, Y: pyrimidine). Any pair of tags created according to these patterns has a guaranteed minimal edit distance of 2; correspondingly these tags are not error correcting in the context of indel and substitution errors. These tags are robust to substitution errors as are predominantly produced by Illumina-type sequencing, requiring four substitutions to erroneously assign any given tag to the wrong pattern.

uvCLAP uses random barcodes that serve as unique molecular identifiers (UMIs) to identify individual crosslinking events from sets of potentially large PCR duplicates. Here, edittag-designed 5’-tags were interleaved with 5 random nucleotides according to the pattern NNNT_1_T_2_T_3_T_4_T_5_NN (N: random nt, T: tag nt), yielding different combinations. The semi-random barcodes positioned adjacent to the 3’-adapter provide additional = 48 random sequences that. In combination, these serve to extend the detectable number of crosslinking events per nucleotide position to at least 49,152 events. To reliably detect the barcodes positioned at both ends of the genomic inserts, uvCLAP by default uses paired-end sequencing. This also allows the use of the genomic positions of both insert ends during PCR duplicate removal to further increase the number of detectable crosslinking events, depending on the number of different insert lengths in the sequenced library.

### uvCLAP data processing

uvCLAP libraries were demultiplexed and adapters removed using Flexbar (version 2.32) (Dodt et al., 2012). Barcodes and UMIs were extracted using custom scripts. To reliably remove readthroughs into barcode regions containing random and semi-random nucleotides, 5 nucleotides (corresponding to the length of semi-random 3’-tags) were clipped from the 3’-ends of first mate reads, 10 nucleotides (corresponding to the length of 5’-tags and UMIs) were clipped from 3’-ends of second mate reads. Since any genomic sequence removed by this step is guaranteed to be contained in the other mate, no information is lost. Bowtie2 (version 2.2.2) (Langmead and Salzberg, 2012) was used to map demultiplexed and processed reads to reference genomes hg19, dm3 and mm10. Uniquely mapped reads were extracted by removing all reads for which multiple alignments could be identified by bowtie2 as indicated by the “XS:i” SAM flag. Alignments sharing UMIs and start coordinates of first and second mate reads were combined into individual crosslinking events. Spurious crosslinking events arising from errors introduced into UMIs were removed as described in (Ilik et al., 2013).

Peaks were called using JAMM (version 1.0.7rev1, parameters “-d y -t paired -b 50 -w 1 -m normal”) (Ibrahim et al., 2015) based on crosslinked nucleotides of biological replicates of foreground and corresponding background libraries.

Pairwise Spearman correlations were calculated using deeptools (version 2.3.5) (Ramírez et al., 2016) based on the numbers of events on the merged peak regions of the full sets of human, fly and mouse experiments.

GraphProt sequence models were trained on 10,000 randomly selected peaks and roughly equal numbers of unbound sequences, using default parameters (GraphProt version 1.1.3) (Maticzka et al., 2014). Unbound sequences were selected by randomly placing peaks within genes with at least one binding site and at least 100 nucleotides apart from any bound site. For training and motif generation, the 60 nucleotides surrounding peak centers were used. Motifs were generated based on the 5% highest-scoring peaks among the 10,000 bound training instances.

Venn diagrams comparing uvCLAP, PAR-CLIP and eCLIP peaks were created using pybedtools (version 0.7.9) (Dale et al., 2011).

The pairwise Spearman correlations and depicted in Figure 2C were calculated using deeptools (version 2.3.5) (Ramírez et al., 2016).

